# Herpes simplex virus 1 inhibits phosphorylation of RNA polymerase II CTD serine-7

**DOI:** 10.1101/2021.06.28.450160

**Authors:** Adam W. Whisnant, Oliver Dyck Dionisi, Arnhild Grothey, Julia M. Rappold, Ana Luiza Marante, Sharada S. Subramanian, Lars Dölken

## Abstract

Transcriptional activity of RNA polymerase II (Pol II) is orchestrated by post-translational modifications of the C-terminal domain (CTD) of the largest Pol II subunit, RPB1. Herpes Simplex Virus type 1 (HSV-1) usurps the cellular transcriptional machinery during lytic infection to efficiently express viral mRNA and shut down host gene expression. The viral immediate-early protein ICP22 interferes with serine 2 phosphorylation (pS2) of the Pol II CTD by targeting CDK9. The functional implications of this are poorly understood. Here, we report that HSV-1 also induces a global loss of serine 7 phosphorylation (pS7). This effect was dependent on the expression of the two viral immediate-early proteins, ICP22 and ICP27. While lytic HSV-1 infection results in efficient Pol II degradation late in infection, we show that pS2/S7 loss precedes the drop in Pol II level. Interestingly, mutation of the RPB1 polyubiquitination site mutation K1268, which prevents proteasomal RPB1 degradation during transcription-coupled DNA repair, displayed loss of pS2/S7 but retained much higher overall RPB1 protein levels even at late times of infection, indicating that this pathway mediates bulk Pol II protein loss late in infection but is not involved in early CTD dysregulation. Using α-amanitin-resistant CTD mutants, we observed differential requirements for Ser2 and Ser7 for production of viral proteins, with Ser2 facilitating viral immediate-early gene expression and Ser7 appearing dispensable. Despite dysregulation of CTD phosphorylation and different requirements for Ser2/7, all CTD modifications tested could be visualized in viral replication compartments by immunofluorescence. These data expand the known means that HSV-1 employs to create pro-viral transcriptional environments at the expense of host responses.

## Introduction

Herpes simplex virus type 1 (HSV-1) is a large, double-stranded DNA virus present in nearly two-thirds of the global population that is the causative agent of the common cold-sore as well as severe skin lesions, life-threatening neonatal encephalitis, and a leading cause of infectious blindness (1). HSV-1 is a paradigm for a virus which induces a profound host shut-off during productive infection by targeting multiple steps of RNA metabolism.

RNA Polymerase (Pol) II is the complex responsible for transcription of all mRNA and several non-coding RNAs. In HSV-1 infection, the virus hijacks Pol II for the transcription of all viral RNAs. Pol II is a multi-subunit complex with the most dynamic regulatory events occurring on the C-terminal domain (CTD) of the largest subunit, RPB1. The RPB1 CTD consists of heptapeptide repeats of the evolutionarily consensus sequence tyrosine-serine-proline-threonine-serine-proline-serine (Y1-S2-P3-T4-S5-P6-S7) (2). Several non-consensus heptapeptide repeats, particularly with variations in the seventh amino acid position, are enriched in the more distal of the 52 mammalian CTD repeats. Each non-proline residue serves as a site of phosphorylation that together help coordinate every step in transcription, with numerous additional post-translational modifications having been described in both the CTD and other regions of RPB1 (3).

Within hours of entering a cell, HSV-1 globally inhibits multiple Pol II processes on host genes such as promoter clearance (4, 5), promoter-proximal pausing (6, 7), and polyadenylation (8) while preserving these functions on viral genes. This selective permissiveness has been attributed to different mechanisms including bulk redirection of DNA-binding proteins to nucleosome-free viral DNA (5, 9) to regulation of specific activities such as mRNA 3’-end formation by the Cleavage and Polyadenylation Specificity Factor (CPSF).

Via its interaction with CPSF, the HSV-1 immediate early protein ICP27 induces the assembly of a dead-end 3’ processing complex, blocking mRNA cleavage. However, 3’-end processing of viral (and a subset of host) transcripts is rescued by the RNA sequence-dependent binding/activator activity of the viral ICP27 protein (10). The viral protein ICP22 has also been shown to interact with Cyclin-dependent kinase 9 (CDK9) (11, 12) and lead to a decrease of phosphorylated CTD Ser2, regarded as a positive marker for transcriptional elongation (13).

Pol II activity is also tightly regulated under conditions of abiotic stress. For instance, a failure of transcription termination has been shown under conditions of hypoxia (14); heat, osmotic, and oxidative stress (15–18); as well as in renal carcinoma (19). Under these conditions, the normal 3’ ends of mRNA are not formed and polymerases transcribe thousands of additional bases downstream of normal termination sites; termed disruption of transcription termination (DoTT), downstream of gene (DoG) transcription, or read-through of polyadenylation (polyA) sites / polyA read-through. Distinguishing which processes are directly regulated by viral proteins and which are disrupted by cellular stress responses becomes imperative to understand transcription during infection.

In this study, we sought to determine HSV-1-induced dysregulation of Pol II CTD modifications and their functional consequences in relation to other stressors. Besides the well-described loss of Ser2 phosphorylation (pS2), we report that HSV-1 infection of primary human fibroblasts also resulted in a global loss of Ser7 phosphorylation (pS7) by 8h post-infection (p.i.), which was not observed in cellular stress responses. Expression of the two viral immediate-early genes ICP22 and ICP27 was necessary to induce loss of pS7 and we provide additional evidence that loss of CTD hyperphosphorylation is separate from bulk RPB1 degradation. Though phosphorylation of both residues is reduced in infection, alanine substitution of Ser7 had no visible impact on viral gene expression while Ser2 substitution was detrimental. Despite Ser7 being dispensable for viral gene expression, its phosphorylation, as well as every other CTD modification examined, could be visualized in viral replication compartments in primary cells. These findings expand the known means of transcriptional regulation by a major human pathogen.

## Results

### Quantification of CTD Modifications During Conditions of polyA Read-through

Considerable overlap exists between genes exhibiting defective polyadenylation during different conditions of osmotic or heat stress and herpes simplex virus infection (20). As transcriptional termination is coordinated in part by CTD phosphorylation, we first determined whether a common disruption of RPB1 modifications occurs under conditions of heat stress (44°C, 2 hours [h]), HSV-1 infection (strain 17syn+, MOI=10, 8h), osmotic stress (80mM KCl, 1h) or oxidative stress induced by 0.5mM sodium arsenite for 1h in primary human foreskin fibroblasts. Timepoints for each condition were based on use in previous studies. Both heat stress and HSV-1 infection induced accumulation of intermediately phosphorylated RPB1 as seen by relative migration in SDS-PAGE (Pol IIi, Fig 1A). While all phospho-serine signals were strongly enriched on hyperphosphorylated Pol II (IIo), the signals of Y1 and T4 phosphorylation and K7 di/tri-methylation more closely correlated to the relative abundance of the different migrating forms of RPB1 in each sample. Interestingly, the phospho-serine signals for heat stress and HSV-1 infection migrated below the hyperphosphorylated IIo signals in mock samples, but remained above the predominate IIi signal for total RPB1 (see E1Z3G in green in S1A Fig).

**Fig 1.**
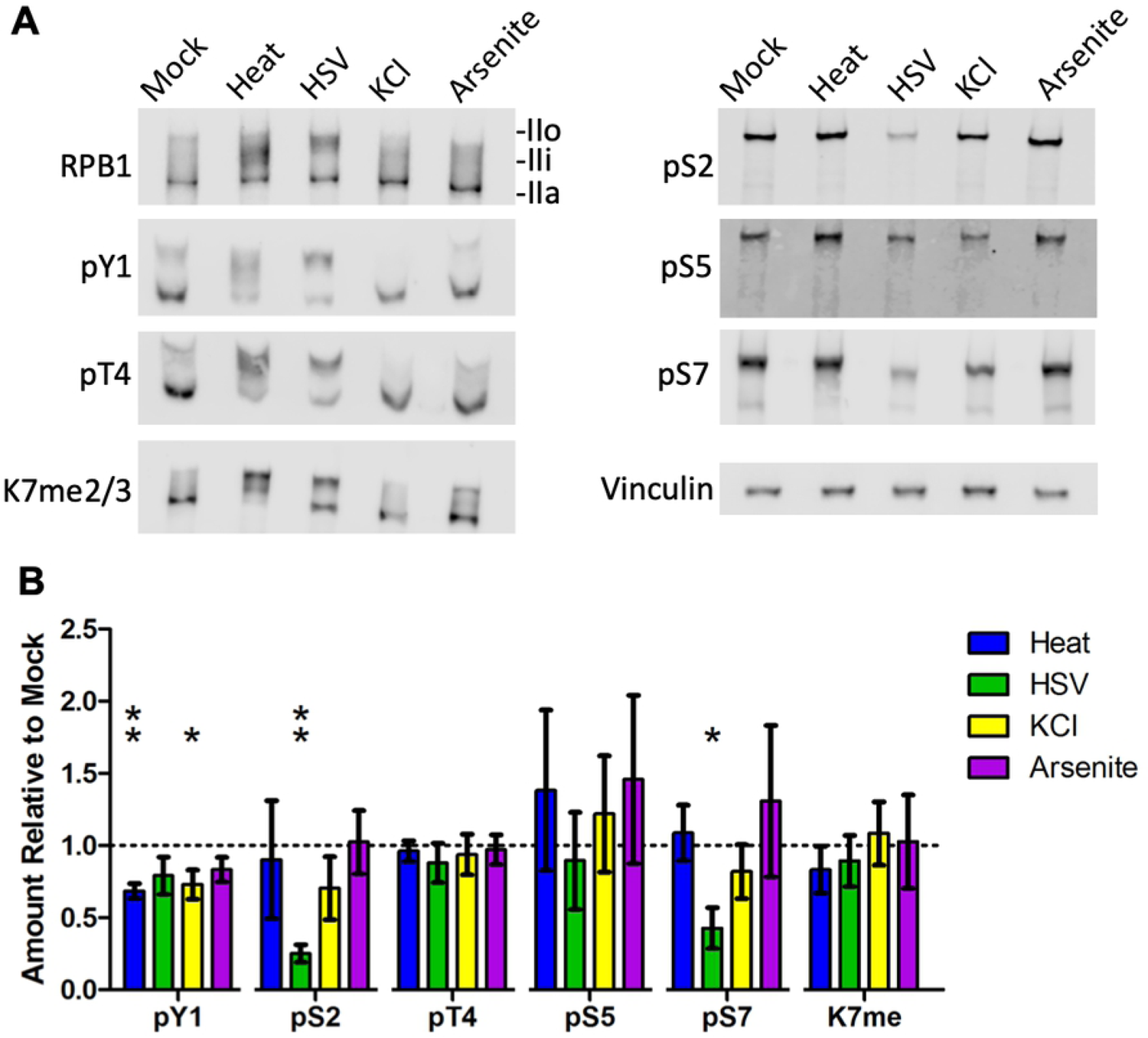
Western blot quantification of RPB1 CTD modifications in primary human fibroblasts during conditions of polyA-site read-through. (A) Human foreskin fibroblasts were subjected to heat (44°C, 2h), HSV-1 infection (strain 17syn+, MOI 10, 8h), osmotic (80mM KCl, 1h), and oxidative (0.5mM NaAsO_2_, 1h) stress and total protein harvested at the end of the stress period. Samples were resolved by SDS-PAGE and probed for levels of RPB1, its CTD modifications, and vinculin as a housekeeping control. Signal for each CTD modification was measured, normalized to the signal for total RPB1 on the same blot, and plotted relative to levels in untreated cells. (B) Mean values of three biological replicates with standard deviations are plotted. Statistically significant differences to mock are indicated as * p < 0.05, ** p < 0.01.

Given the various migration patterns across all samples, we thus quantified signals for CTD modifications across the entire range of RPB1 bands and normalized to total RPB1 signals on the same blot. For all stress conditions, a slight (20-30% on average) reduction in tyrosine 1 phosphorylation (pT1) was observed relative to mock (Fig 1B). Surprisingly, threonine 4 phosphorylation (pT4), which predominately accumulates around transcription termination sites and is involved in 3’-end formation (21, 22), remained unaffected in all conditions; as did serine 5 phosphorylation (pS5) and K7 di/tri-methylation levels (K7me). HSV-1 infection was the only condition observed to heavily impact CTD modifications, causing a ∼70% reduction in pS2, as previously described, and a ∼50% loss of serine 7 phosphorylation (pS7).

As previous studies have shown binding of monoclonal antibodies to the CTD can be affected by the identity and phosphorylation of nearby amino acids (23–26), we tested a panel of different CTD phospho-serine antibodies for each serine residue on the same samples. The data for CTD phospho-serines in Fig 1 are for selected antibodies whose epitope biases have been evaluated by in other studies (23, 25), with results for all antibodies in S1 Fig. Of five antibodies tested for pS2, all showed a reduction of pS2 in HSV-1 infection, while three antibodies showed a ∼2-fold reduction for heat and two for salt stress (S1A Fig). Previous studies have observed similar gel migration shifts after heat shock in mammalian and *Drosophila* cells (27–29), with a decrease of pS2 and pS5 reported in mouse cells (30). One of three antibodies showed increases of pS5 levels for heat stress and oxidative stress, while one of two antibodies showed a reduction of pS7 in all conditions (S1 Fig). Overall, these data indicate that there that HSV-1 causes a loss of two major CTD modifications while there is no major CTD modification indicative of general stress responses beyond a slight reduction of pY1. Dephosphorylation of Y1 is directly involved in recruiting termination factors in yeast (31), and mammalian RPB1 mutants where the distal 3/4 of Y1’s are substituted for phenylalanine also exhibit termination defects (32). However, it remains unclear whether a ∼20% global reduction in pY1 would be sufficient to recapitulate this phenotype during cellular stress.

### Loss of CTD serine 7 phosphorylation in HSV-1 infection is due to expression of ICP22 and ICP27

The loss of pS2 during HSV-1 infection has been linked to inhibition of CDK9 by the viral immediate-early (IE) protein ICP22 (11). As CDK9 can also phosphorylate Ser7 (3), we investigated the role of ICP22 and other viral gene products in the loss of pS7. A time course of infection, with and without the DNA replication inhibitor phosphonoacetic acid (PAA) to prevent viral late gene expression, demonstrated that viral late gene products are not required for the observed pS7 loss (Fig 2A,B), with a steady decrease beginning at 4h p.i. To specifically test immediate-early (IE) genes, a cycloheximide reversal assay was performed. In this assay, cells are infected and then treated with cycloheximide to allow transcription – but not translation – of the five viral IE genes, which do not require synthesis of new viral proteins for expression. After 4h, cycloheximide was removed and replaced with actinomycin D for an additional 8h to allow translation of the accumulated viral IE mRNA but prevent transcription of early and late genes. In these conditions however, a noticeable decrease in all forms of Pol II due to degradation was observed (S2A Fig). This is in line with previous reports using actinomycin D suggesting that the normal virally induced CTD modifications or even RPB1 stability require ongoing transcription (33).

**Fig 2.**
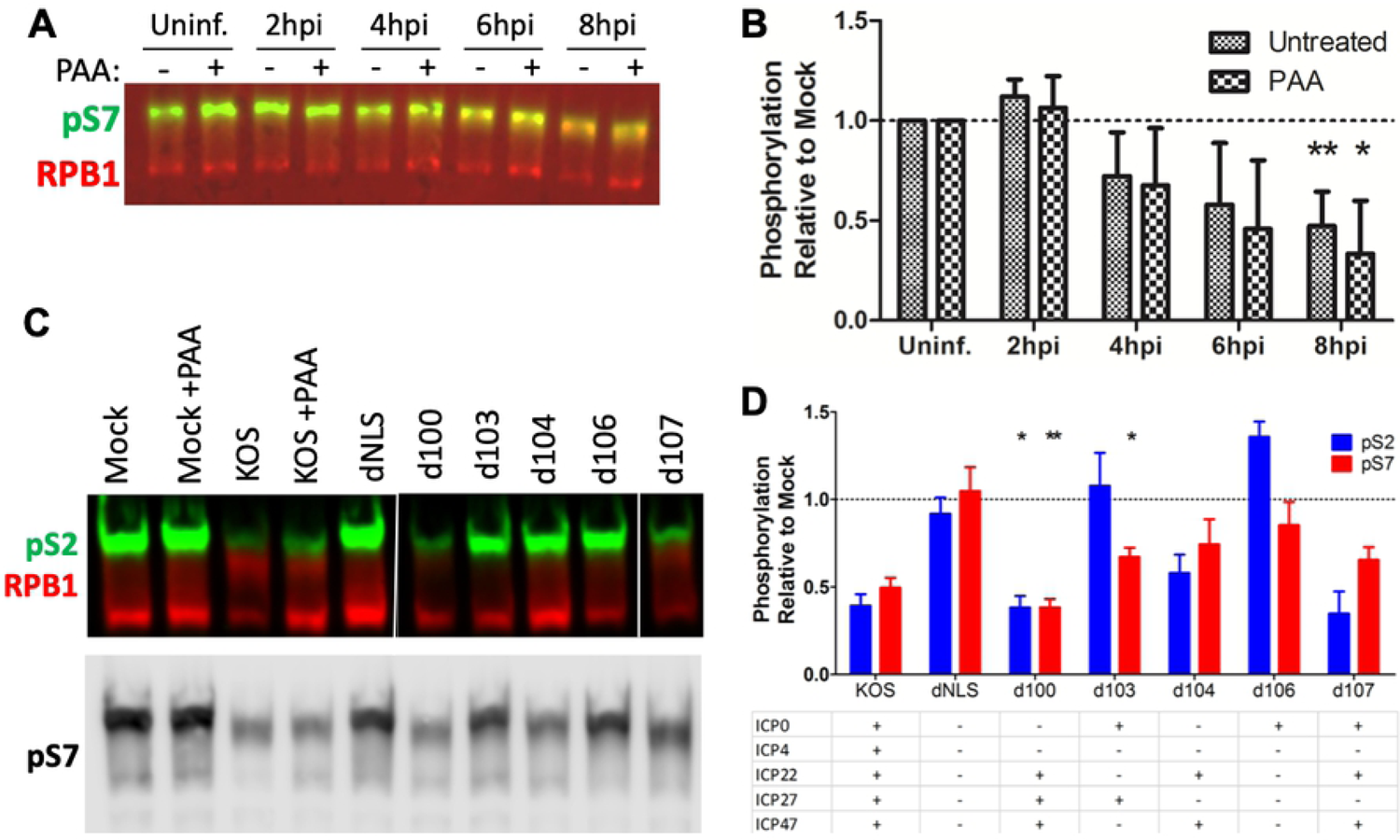
Loss of CTD serine 7 phosphorylation in HSV-1 infection requires immediate-early proteins ICP22 and ICP27. (A) Human foreskin fibroblasts (HFF) were infected with HSV-1 17syn+ with and without DNA replication inhibitor phosphonoacetic acid (PAA) and total protein harvested at given timepoints. Levels of CTD serine 7 phosphorylation (pS7) relative to total RPB1 were quantified by Western blotting. (B) Means of at least three replicates relative to the corresponding mock-infected sample plotted with standard deviations. (C) HFF cells infected with HSV-1 KOS treated with PAA or KOS-derived mutants lacking the combinations of immediate-early genes indicated in the table in the bottom right. White bars indicate removal of irrelevant samples. Total protein was harvested at 8h p.i. and pS2/7 levels determined by Western blot. (D) Quantification of phospho-serine levels normalized to total RPB1. Means of three replicates normalized to mock-infected samples with PAA with standard error are plotted, values from KOS are from PAA-treated samples. Statistically significant differences to mock are indicated as * p < 0.05, ** p < 0.01.

Of the five viral IE genes, four (ICP0, ICP4, ICP22, and ICP27) show nuclear localization and viral mutants lacking each of these genes were tested for their impacts on pS7. Contrary to pS2, which accumulated to higher levels without ICP22 compared to the wild-type strain, no individual viral IE gene seemed sufficient to account for the loss of pS7 (S2B Fig). The addition of PAA with these mutants prevented some of the pS7 loss, indicating that viral late genes and/or DNA replication, though not required, can facilitate CTD remodeling as previously described (12,13,34). Importantly, no loss of pS7 was observed upon infection with a mutant virus whose genome is not trafficked to the nucleus due to the removal of the nuclear localization signal in the UL36 protein (dNLS, Fig 2C,D). Therefore, viral tegument proteins delivered by the incoming virus particles are not sufficient to induce pS7 loss, as indicated by the retention of pS7 when using cycloheximide alone (Fig. S2A).

As our data suggested the involvement of more than one viral gene product, a panel of viruses lacking the major viral transcription factor ICP4 and combinations of the other IE genes were tested. It is important to note that viral gene expression by these viruses is restricted to the remaining four immediate early genes (ICP0, ICP22, ICP27 and ICP47.5). The strongest reduction (∼50%) in pS7 was observed in viruses expressing both ICP22 and ICP27 (KOS and d100, Fig 2C,D). Viruses lacking either ICP22 or ICP27 had an intermediate (∼25%) loss of pS7, while a virus lacking both proteins (d106) had no significant reduction. The contribution of ICP22 to the loss of both pS2 and pS7 can be easily attributed to inhibition of CDK9, while ICP27’s impact on pS7 is less readily apparent. ICP27 is known to co-precipitate with RPB1 (35, 36) and other transcription factors such as SPT5; the latter interaction facilitated by CDK9 activity (37). ICP27 has also been previously reported to reduce Ser2 phosphorylation in HeLa and RSF (36, 38) –but not Vero (13)– cells. However, as the other immediate-early mRNAs exhibit less cytoplasmic accumulation without ICP27 (39), an effect on ICP22 expression cannot be ruled out for the impact on pS2. These data indicate that while loss of ICP27 did not alter ICP22-mediated effects on serine 2 phosphorylation in infected primary human fibroblasts, ICP22 and ICP27 work in a complementary manner to deplete serine 7 phosphorylation that does not require viral late gene expression.

### HSV-1 utilizes multiple proteasome-dependent pathways to regulate RPB1

Early studies have shown that total transcriptional activity of all RNA polymerases in HSV-infected cells rapidly shuts down after the peak of viral gene expression around 8h p.i. (40–42), associated with Pol II holoenzyme remodeling (43) and degradation of RPB1 itself at later timepoints (33). In addition to viral replication compartments, RPB1 localizes to virus-induced chaperone (VICE) domains (36, 38) which have been proposed to serve as sites of quality control for protein folding and/or complex assembly as they contain numerous chaperones and proteasome components (44, 45). Protease activity and different types of mono- and polyubiquitination have been detected in VICE domains, though it is not clear if all proteins trafficked to these compartments are fated for degradation or if these can serve as assembly sites for complicated higher-order protein structures (46).

While the exact roles of VICE domains in regulating transcription are unknown, proteasome inhibitors which block VICE domain formation (36, 38) were previously found to prevent the loss of pS2 (36), and we observed this for pS7 as well (Fig 3A). Efficacy of the proteasome inhibitors could be visualized by increased p53 levels and laddering indicating polyubiquitination, and for RPB1 specifically accumulation of a high molecular weight (MW) band likely corresponding to ubiquitinated RPB1 (38) and lower MW bands indicative of degradation products could be observed in both infected and uninfected cells (S3A Fig). Interestingly, the migratory shift of RPB1 from IIo to IIi during gel electrophoresis could still be observed in infected cells even though treated samples had relatively equal levels of pS2 and pS7 as uninfected controls (S3B Fig). We are unable to measure the levels of ICP22 expression under these settings due to the lack of a commercial antibody, but previous work demonstrated that ICP22 expression is not altered by proteasome inhibition (47).

**Fig 3.**
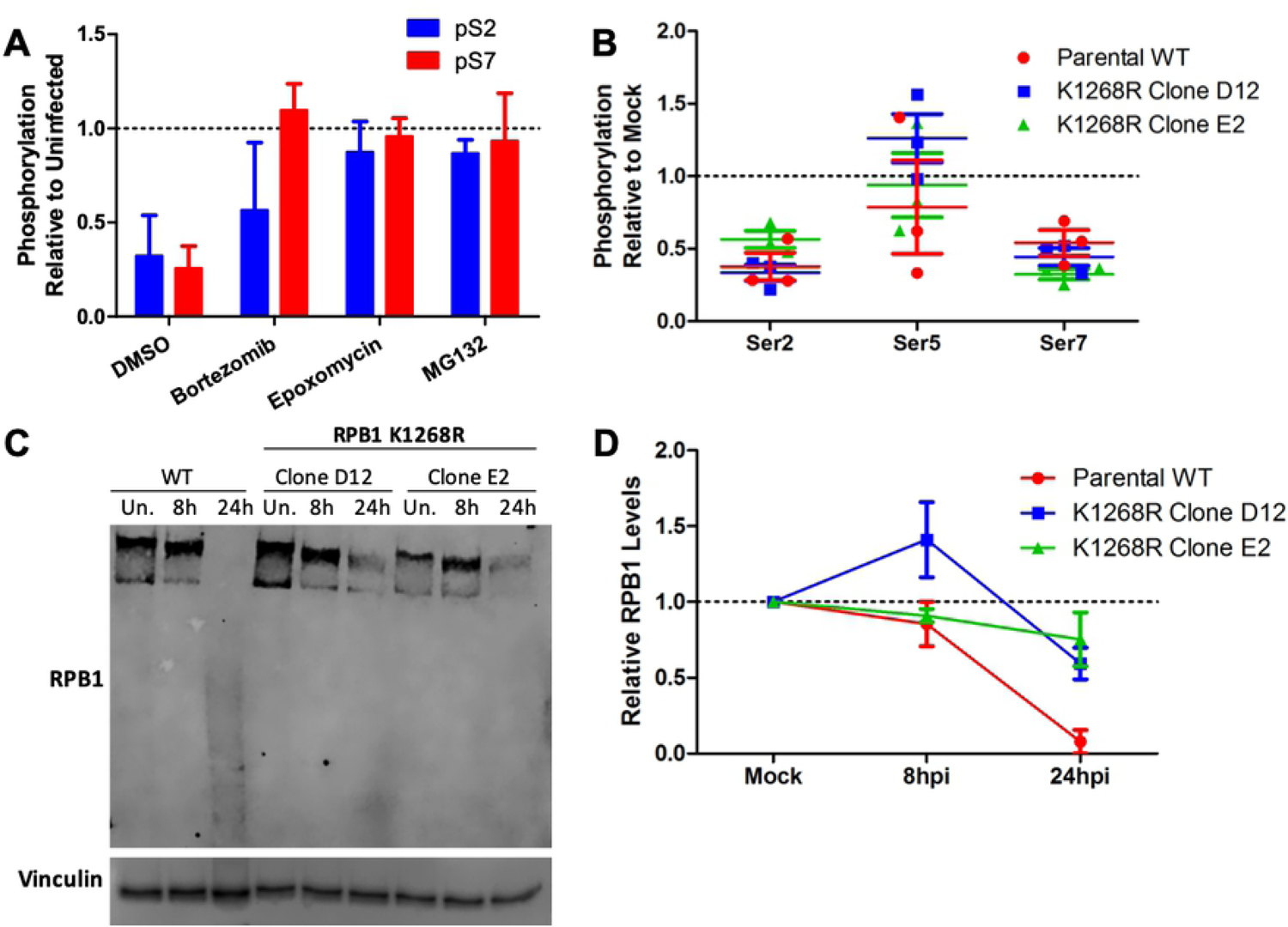
Effects of proteasome inhibitors and RPB1 polyubiquitination mutants on HSV-1-induced RPB1 remodeling. (A) Human foreskin fibroblasts were infected with HSV-1 17syn+ and treated with indicated proteasome inhibitors for 8h before collection of total protein and quantification of CTD Ser2 and Ser7 phosphorylation by Western blotting. Plotted are means of two replicates with standard deviations. (B-D) 293T parental wild-type (WT) and clonally derived RPB1 K1268R mutant cell lines were infected with HSV1(17+)-LoxCheVP26 and total RPB1 or phosphorylated CTD serine residues quantified by Western blotting. (B) Relative levels of CTD serine phosphorylation at 8h p.i.; plotted are three individual replicates with longest lines indicating means and shortest standard error ranges. (C) Representative Western blot of RPB1 levels in uninfected (Un.) 293T WT and RPB1 K1268R or cells infected for 8 and 24h. (D) Relative RPB1 levels normalized to vinculin in infected 293T WT and RPB1 K1268R cells; plotted are means of at least three replicates with standard error.

Rice and colleagues have suggested that ICP22-mediated inhibition of pS2 is mechanistically distinct from the degradation of hyperphosphorylated RPB1 (13). To this point, we observed no significant loss of total RPB1 levels by 8h p.i. or immediately following heat, osmotic, or oxidative stress using five different RPB1 antibodies (S4 Fig), indicating CTD remodeling as the major effector or degradation of a very small percentage of RPB1 molecules (as detectable by Western blot) which contain the vast majority of Ser2/7 –but not Ser5– phosphorylation in the cell.

The most well-studied RPB1 degradation pathway occurs during transcription-coupled repair where elongating Pol II stalls at sites of DNA damage and signals to repair machinery via polyubiquitination of RPB1, leading to release of RPB1 from DNA and subsequent degradation. Recent work demonstrated that this polyubiquitination occurs on a single lysine, K1268; and while other sites of monoubiquitination exist, mutation of this single lysine to arginine completely prevented degradation of RPB1 after UV irradiation (48). Two monoclonal cell lines bearing the RPB1 K1268R mutation exhibited an equal loss of serine 2 and 7 phosphorylation during HSV-1 infection as the parental wild-type (WT) cell line by 8 h p.i. (Fig 3B, S5A). As in primary fibroblasts, no major loss of RPB1 was observed by 8h p.i. in any of the cell lines. By 24h p.i. however, RPB1 in WT cells was hardly detectable; migrating almost exclusively as a lower MW smear while RPB1 K1268R mutant cells exhibited a ∼50% retention in total RPB1 protein (Fig 3C,D). Visualization of mCherry fused to the viral late gene VP26 indicated that viral gene expression at 8 and 24h was comparable across all cell lines and that this effect was not due to global differences in viral gene expression (S5B Fig). This demonstrates that the RPB1 ubiquitination pathway active during DNA damage is not involved in CTD remodeling during the bulk of viral gene expression, but may degrade Pol II at later times during virion assembly and genome packaging. It is possible that the 50% reduction of RPB1 in the K1268R cells may not be due to degradation, but standard protein turnover coupled with the virus-induced shutoff of host mRNA translation as RPB1 is reported to have a half-life of 6h in mammalian cell lines (49). Virus production in these cells was measured to determine if RPB1 degradation facilitates productive infection and virus release, however the two clones bearing the K1268R mutation gave differential phenotypes as the mutant clone D12 produced viral progeny equal to WT cells while the E2 clone was consistently ∼5-fold lower than the other two cell lines (S5C Fig). Overall, these data indicate that while proteasome-dependent pathways are active in CTD remodeling during lytic HSV-1 infection, they are separate from transcription-coupled repair and precede the bulk degradation of RPB1.

### Herpesviral DNA recruits, but does not require, all CTD modifications

During infection, Pol II is strongly enriched in viral replication compartments (9): large compartments predominately assembled by viral DNA and the viral transcription factor ICP4 which recruit numerous factors for transcription, DNA replication, and virion assembly while excluding host proteins such as histones and gene silencing factors (50). We sought to determine which CTD modifications localized to viral DNA by immunofluorescence. Even though HSV-1 infection reduced global pS2 and pS7 levels, these modifications were still strongly enriched in ICP4 foci, as were all other CTD modifications tested (Fig 4); indicating that no major CTD mark is excluded from the viral genome.

**Fig 4.**
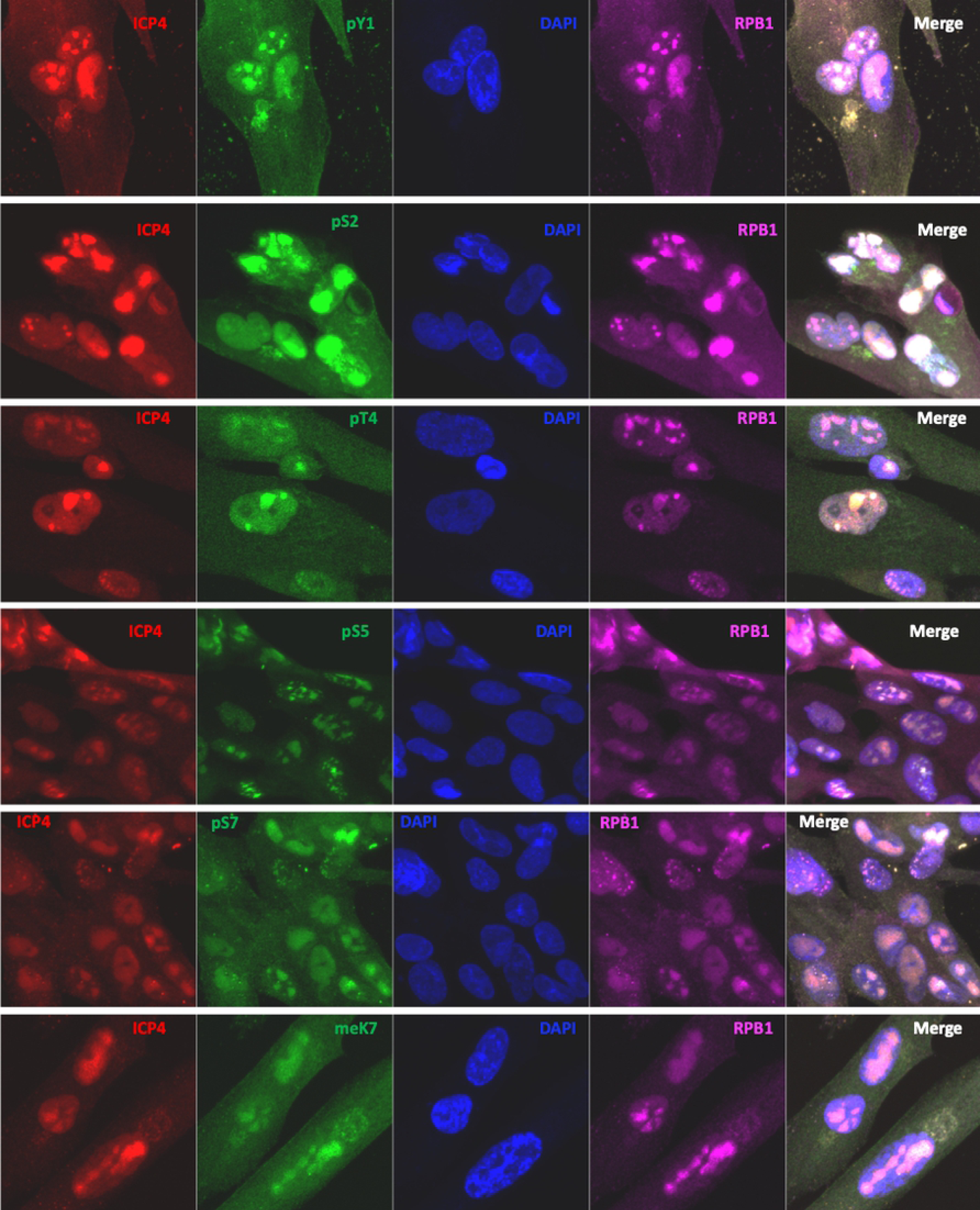
Co-localization of RPB1 CTD modifications with viral replication compartments. Human foreskin fibroblasts were infected with HSV-1 strain 17syn+ for 8h before fixation. Antibody staining was performed for viral protein ICP4, RPB1, and its respective CTD modifications. p=phosphorylation, me=methylation.

Previous studies have shown that chemical inhibition of CDK9 reduces herpesviral gene expression (51, 52), indicating that HSV-1 still requires some degree of phosphorylation on the CTD and/or other transcription factors. We decided to investigate the requirements of viral genes for CTD serines 2 and 7 using an amanitin-based RPB1 replacement assay. HEK-293T cells were co-transfected with a plasmid expressing the blue fluorescent protein Azurite as a visual control for transfection efficiency and vectors expressing α-amanitin-resistant, HA-tagged RPB1 constructs (53, 54) containing either a wild-type (WT) CTD or mutant CTDs. The mutations are non-phospho-accepting alanine substitutions in place of serines 2 or 7 (S2A and S7A, respectively) or the phospho-serine mimic glutamic acid at serine 7 (S7E). After 24h, α-amanitin was added to degrade endogenous RPB1 for an additional 24h, followed by infection with fluorescent HSV-1 expressing eYFP-ICP0 as an immediate-early gene marker and gC-mCherry as a late gene marker (55). As shown in Fig 5, detectable levels of IE and late genes could be visualized in cells with replaced WT CTD at 24h p.i., albeit at substantially lower levels than amanitin-untreated cells, but not with empty pcDNA3 vector control. As expected, both early and late viral gene expression was reduced with CTD S2A mutants. In contrast, S7A mutations expressed both classes viral genes equally well as the WT CTD. Interestingly, the phospho-mimic S7E mutations completely ablated viral gene expression, moreso than S2A, though this is likely due to reduced recruitment and/or 3’-end processing (54) as even Azurite expression was reduced. Western blotting for the plasmid-expressed, HA-tagged RPB1 verified that expression of the exogenous RPB1 mutants were comparable across conditions (S6 Fig). These data demonstrate that while viral DNA localizes all major CTD modifications, not all are required for viral late gene expression. In the case of CTD serine 7, the amino acid itself appears completely dispensable during lytic infection.

**Fig 5.**
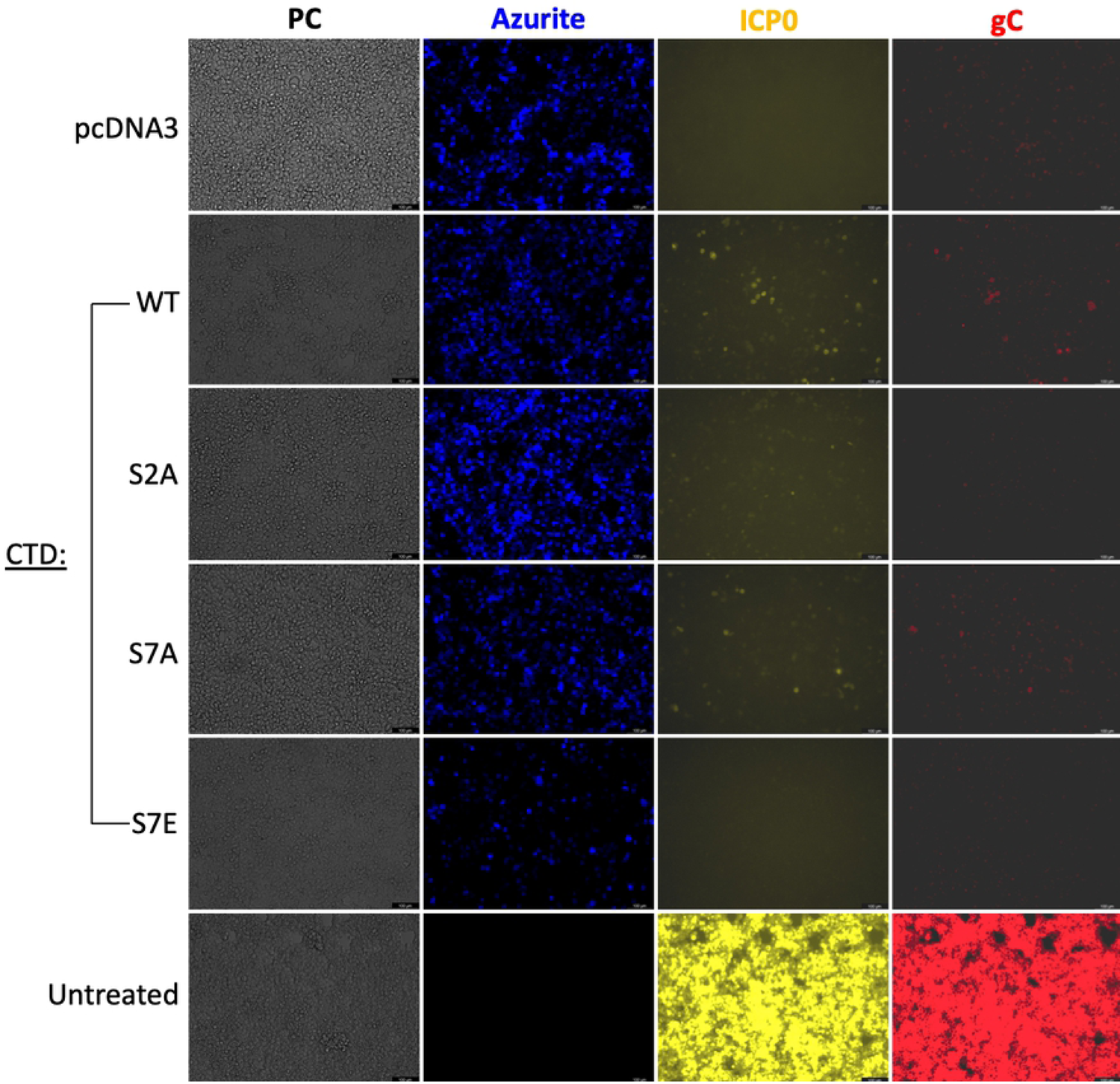
CTD serine 2, but not serine 7, is required for efficient viral gene expression. HEK-293T cells were co-transfected with a construct expressing Azurite blue fluorescent protein as a transfection marker and plasmids expressing amanitin-resistant RPB1 with wild-type (WT) or indicated mutant CTDs, or the control vector pcDNA3 lacking any polymerase. The next day, α-amanitin was added to degrade endogenous RPB1 for 24h followed by infection with an eYFP-ICP0-/gC-mCherry-expressing HSV-1. Live-cell imaging was performed at 24h p.i. All images were acquired with the same settings, with infection of untransfected, amanitin-untreated cells included as a reference for the relative level of viral gene expression. PC: phase contrast, S2A: serine 2 to alanine, S7A: serine 7 to alanine, S7E: serine 7 to glutamate. Scale bars indicate 100μm.

## Discussion

Post-translational modification, particularly phosphorylation, of the RPB1 CTD is at the core of numerous transcriptional responses. Release of CDK9 from the 7SK snRNP and resulting transcription P-TEFb-dependent genes greatly impacts mammalian cell survival during genotoxic stress (56) and interferon responses during viral infection (57). As phosphorylation of multiple CTD residues (namely Tyr1, Ser2, and Thr4) are involved in termination of mammalian protein-coding genes, we reasoned that there could be a core dysregulation of the CTD code in response to stress. However, we observed no major changes in the CTD shared across all conditions tested. Both HSV-1 infection and heat stress induced the intermediate-migrating IIi form of RPB1, while only HSV-1 showed a consistent loss of pS2 and pS7 with all antibodies tested. To avoid antibody bias, future work should implement mass spectrometry-based assays (2, 58) to quantify the phosphorylation states of the IIi forms of RPB1 in heat stress and HSV-1 infection. Though different challenges to cellular homeostasis can have similar global impacts on gene expression, particularly in regards to inhibiting polyadenylation, our data demonstrate that this is not due to a common, global disruption of the RPB1 CTD.

Western blotting of different viral mutants implicates *de novo* expression of both ICP22 and ICP27 in the loss of pS7. While ICP22’s known ability to regulate CDK9 can account for this protein’s contribution, ICP27 may work by other means. Beyond simply supporting expression of other viral genes, ICP27 binds directly to RPB1 and could disrupt downstream interactions and phosphorylation. Both ICP22 and ICP27 have been implicated in VICE domain formation, and it is possible that RPB1 may interact with proteins in these domains that limit its ability to be phosphorylated. Though VICE domains contain numerous components involved in protein misfolding and degradation, we favor the view that they facilitate RPB1 remodeling and trafficking over proteolysis as we do not observe a major decrease in RPB1 levels until well after the decrease in pS2/7.

Our data supports previous observations that ongoing viral transcription is required for CTD remodeling, as even relatively weak expression of immediate early genes can trigger RPB1 degradation in cycloheximide/actinomycin D reversal assays. This particular degradation pathway is likely separate from the one during DNA damage as it occurs when transcription is chemically inhibited. We observed that degradation of RPB1 mediated by polyubiquitination of K1268 is not involved in the loss of pS2/7, but stabilizes RPB1 24h p.i. This suggests that there are multiple degradation pathways triggered by HSV-1 at different stages of infection and reside in a balance with ongoing viral transcription or trafficking between VICE domains and replication compartments (RCs). Degrading Pol II on viral DNA may preserve genomes in settings of abortive infection by halting the production of viral antigens or innate immune triggers like double-stranded RNA, or help maintain latency with leaky transcription. It will be of interest to experimentally determine ubiquitination sites at early and late timepoints of HSV-1 infection by mass spectrometry to better understand how the proteasomal machinery is involved with viral transcription cycles and shuttling of RPB1 between the various nuclear compartments.

The interaction of ICP22 and CDK9 has been proposed to benefit HSV-1 by causing reduced transcriptional elongation rates of cellular mRNA as a means of countering host responses (11), and even to regulate viral genes themselves (59). However, chemical inhibition of CDK9 also reduces viral gene expression and our data with S2A mutants recapitulates this phenotype, indicating a beneficial role for at least intermediate pS2 levels in regards to viral genes. Contrary to this, polymerases with S7A mutations expressed viral proteins comparable to the WT CTD, demonstrating that the entire CTD code is not required for herpesviral gene expression despite the fact that all modifications we could test are localized to viral replication compartments. The presence of pS7 on the viral genome may therefore be a consequence of recruiting kinases for their other, necessary phosphorylation events or serve to sequester transcription factors away from host chromatin. We also cannot rule out the possibility that Ser7 regulation may be more impactful during reactivation from latency over lytic infection or during inflammatory responses. Indeed, inhibition of another Ser7 kinase, CDK7, has been shown to reduce cytokine release and inflammation (60, 61), but more work is required to determine contributions from Ser7 vs. other CDK7 targets, especially CTD Ser5.

Currently, the best described role of pS7 is recruitment of the Integrator complex to snRNA genes (62) for 3’-end formation of Pol II-derived snRNA (63). Disrupting snRNA formation by itself is unlikely to influence lytic infection as the rapid redirection of Pol II activity to viral genomes and transcriptional shutdown within the span of 8-12h would not be expected to alter the abundant pool of mature splicing factors, and most viral genes are unspliced late in infection when these termination defects are the greatest. Integrator also has direct roles in regulating protein-coding genes, and knockdown of its catalytic subunit recapitulates termination defects on a subset of host mRNAs disrupted during osmotic stress (64), though our previous work implicates ICP27-CPSF interactions as the primary determinant during HSV-1 infection (10). Work is ongoing to analyze the role of integrator in HSV-1 infection. The reduction of pS7 could also result in more subtle, global consequences on host gene expression as pS7 has been proposed to suppress cryptic transcription (65) as well as stimulate Ser2 phosphorylation by CDK9 and CDK12 (66, 67). Our future efforts will involve mapping the genomic locations of pS/7 during infection by ChIP/mNET-seq to better understand the impacts of herpesviral-induced CTD remodeling.

## Materials and Methods

### Cell lines and viruses

Human foreskin fibroblasts were purchased from ECACC and ATCC. BHK-21 cells were purchased from ATCC. 293T RPB1 K1268R clones D12 and E2 as well as the corresponding parental cell line (48) were gifts from Jesper Svejstrup, Francis Crick Institute. Vero cells were provided by Beate Sodeik, Hannover Medical School, and Vero derivatives E5 (68) and F06 (69) were gifts from Neal A. DeLuca, University of Pittsburg and Vero 2-2 (70) from Rozanne M. Sandri-Goldin, UC Irvine. RSC and RSC-HAUL36 (71) were provided by Peter O’Hare, Imperial College London. All cells were cultured in Dulbecco’s Modified Eagle Medium (DMEM), high glucose, pyruvate (ThermoFisher #41966052) supplemented with 1x MEM Non-Essential Amino Acids (ThermoFisher #11140050), 1mM additional sodium pyruvate (ThermoFisher #11360070), 10% (v/v) Fetal Bovine Serum (FBS, Biochrom #S 0115), 200 IU/mL, penicillin and 200 µg/mL streptomycin. All cells were incubated at 37°C in a 5% (v/v) CO_2_-enriched incubator.

KOS strain ICP4 mutant n12 (68) and KOS 1.1 ICP27 mutant 27-lacZ (70) were propagated on E5 cells and Vero2-2 cells, respectively. KOS BAC-derived VP1-2ΔNLS (72) was cultured as previously described in RSC cells. Wild type strains 17syn+ and F, along with BAC-derived strains KOS and HSV1(17+)-LoxCheVP26 (73) were propagated on BHK-21 cells as well strain 17 ICP0 mutant dl1403 (74), strain F ICP22 mutant Δ325 (75), as well as the KOS eYFP-ICP0/gC-mCherry BAC-derived virus (55), which was generously provided by Colin Crump, University of Cambridge. KOS mutants d100, d103, d104, d106, and d107 (76) were grown and titered on F06 cells.

293T cell lines were seeded onto poly-lysine-coated dishes for infection experiments. All infections, except those for virus growth curves, were performed with an MOI of 10 by diluting virus in serum-free DMEM and adsorbing for 1h at 37°C to cells plated the previous day. Virus production assays were similarly infected but with an MOI of 0.01. Afterwards, inoculum was replaced with culturing medium or medium containing concentrations of compounds described below.

### Stress and drug treatments

Heat stress was initiated by adding pre-warmed 44°C medium to cells and culturing for 2h at 44°C. Salt stress and oxidative stress were initiated by adding KCl or sodium arsenite to concentrations of 80mM and 0.5mM, respectively, for 1h. Mock cells were harvested at timepoint 0 and infected cells at 8h p.i. Phosphonoacetic acid (PAA, Sigma #284270) was used at a concentration of 300 µg/mL when indicated. Bortezomib (Selleckchem #S1013) was used at a concentration of 5µM, epoxomycin (Cayman Chemical #10007806) and MG132 (Sigma #M7449) both at 10µM, with equal volumes of DMSO vehicle used as control. Actinomycin D was used at 5 µg/mL and cycloheximide at 100 µg/mL for the times indicated in the text. Uninfected cells for drug treatments were harvested at the final timepoints as infected samples.

### Western blotting

A full description of antibodies is found in S1 Table. Total protein samples were harvested by lysing cells directly in Laemmli buffer at given timepoints. Samples were resolved on 6% tris-glycine SDS PAGE gels with a 4% stacking gel, both 37.5:1 acrylamide:bisacrylamide, and transferred overnight in tris-glycine buffer containing 20% (v/v) methanol to 0.45µm nitrocellulose membranes. Membranes were rinsed with deionized water and blocked in tris-buffered saline with 0.1% (v/v) Tween 20 (TBST) with 5% (w/v) milk for 1h before overnight binding with primary antibodies diluted in blocking buffer. Samples were washed in TBST, blocked for one additional hour, and secondary antibodies allowed to bind for one hour before final TBST washes and visualization on a LI-COR Odyssey Fc imaging system. Deviations from this procedure are indicated in the relevant figure legends.

Band densitometry was performed using Image Studio Light (LI-COR). Total RPB1 levels for each sample were normalized to vinculin signal on the same membrane. Signals for each CTD modification were normalized to total RPB1 levels on the same membrane. The relative ratios are compared to mock samples on the same membrane, which was set to 100%. Statistical significance was determined using two-way ANOVA in GraphPad Prism.

### Immunofluorescence

3×10^4^ HFF cells were seeded onto 8 well glass chamber slides (Ibidi #80841). The following day, cells were infected at an MOI of 10 and fixed 8h p.i. in 4% formaldehyde in PBS for 15 minutes. After washing in PBS, cells were incubated in permeabilization buffer (10% [v/v] FBS, 0.5% [v/v] Triton X-100, 250mM glycine, 1x PBS) for 10 minutes, then blocked for 1h in blocking buffer (10% [v/v] FBS, 250mM glycine, 1x PBS). Primary antibodies were incubated overnight at 4°C in 10% (v/v) FBS and 1x TBS. The secondary antibodies were incubated in 10% FBS in 1x TBS for 1h at room temperature with 1 μg/mL 4’,6-diamidino-2-phenylindole (DAPI). A full description of antibodies is found in S1 Table. Coverslips were washed in water before mounting them in medium containing Mowiol 4-88 and 2.5% (w/v) 1,4-diazabicyclo[2.2.2]octane (DABCO). All steps before antibody binding were followed by three 5-minute washes in PBS, while TBS was used after antibody binding. Samples were imaged on a Zeiss LSM 780 where slices of 0.5µm were taken and intensities of each slice summed in Fiji (77) to produce the 2D images provided. Live-cell imaging of fluorescent viruses was performed on a Leica DMi8.

### RPB1 replacement assay

5×10^4^ HEK-293T cells were seeded in poly-lysine-coated 48-well plates. The next day, cells were transfected with 15ng of pLV-Azurite (gift of Pantelis Tsoulfas, Addgene #36086) and 250ng of vectors expressing HA-tagged, amanitin-resistant RPB1 constructs with the mutant CTDs or pcDNA3 as a negative control via polyethylenimine (PEI). The WT and S2A mutant are described in (53), the S7A and S7E mutants in (54). All RPB1 constructs were generous gifts of Dirk Eick, LMU Munich. After 24h, media was exchanged for fresh growth medium with 2.5 μg/mL α-amanitin for an additional 24h. Cells were infected with eYFP-ICP0/gC-mCherry HSV-1 at MOI 10 as described above in serum-free medium for 1h. This inoculum was removed and amanitin-containing growth medium reapplied until imaging 24h p.i.

## Acknowledgements

We would like to thank Dirk Eick for the mutant RPB1 vectors and methyl-CTD K7 antibody, Neal DeLuca and Roz Sandri-Goldin for providing HSV-1 deletion mutants and complementing cell lines, as well as Colin Crump and Beate Sodeik for fluorescent BAC viruses. This work was supported by the Deutsche Forschungsgemeinschaft (DFG grant DO1275/6-1) and the European Union (ERC-2016-CoG 721016-HERPES) to L.D. A.W.W. was the recipient of a generous grant from the Alexander von Humboldt Foundation and the German Federal Foreign Office.

## Contributions

A.W.W. and L.D. conceived and designed the experiments and wrote the paper. A.W.W. performed the experiments with the help of O.D.D., A.G., J.M.R., A.L.M., and S.S.S.

## Supporting Information

**S1 Fig. Measurement of RPB1 CTD serine phosphorylation after exposure to stress using different antibodies.** Human foreskin fibroblasts were subjected to mock treatment (lane 1), heat stress (lane 2; 44°C, 2h), HSV-1 infection (lane 3; strain 17syn+, MOI 10, 8h p.i.), osmotic (lane 4; 80mM KCl, 1h), and arsenite (lane 5; 0.5mM NaAsO_2_, 1h) treatment and total protein harvested at the end of the stress period. Quantification of CTD serine 2 (**A**), serine 5 (**B**), and serine 7 (**C**) phosphorylation was performed using the listed antibodies via Western blotting. Phospho-serine levels were normalized to total RPB1 levels and the relative ratio compared to mock-treated samples on the same membrane. Plotted are the means of three biological replicates with standard error with representative Western blots on the right. Statistically significant differences to mock are indicated as * p < 0.05, ** p < 0.01, *** p < 0.001.

**S2 Fig. CTD serine 7 phosphorylation in cycloheximide reversal and infection with HSV-1 immediate-early gene mutants.** (A) Human foreskin fibroblast (HFF) cells infected with HSV-1 17syn+ and treated with DMSO, actinomycin D (ActD), cycloheximide (CHX), or cycloheximide reversal (SWAP; 4h CHX followed by 8h ActD) and total protein analysed at 12h p.i. for levels of CTD Ser7 phosphorylation (pS7) and each indicated protein. Samples for total RPB1 were resolved on 3-8% Tris-Acetate gels to better capture lower molecular weight degradation products. (B) HFF cells infected with indicated HSV-1 mutants at MOI 10 for 1h, and inoculum replaced with growth media or media containing phosphonoacetic acid (PAA) for 8h. Total protein was harvested and levels of pS2, pS7, and RPB1 analyzed by Western blotting. Wild-type (WT) strains F, KOS, and 17syn+ are shown next to corresponding mutants for ICP22 (Δ22), ICP4 (Δ4), ICP27 (Δ27), and ICP0 (Δ0). (C) Quantification of Western blots described in (B). Means of at least three biological replicates for PAA-treated samples are plotted with standard errors. Statistically significant differences to mock are indicated as * p < 0.05, ** p < 0.01.

**S3 Fig. Proteasome inhibition blocks the loss of CTD serine 2 and serine 7 phosphorylation during HSV-1 infection.** Human foreskin fibroblasts were infected with HSV-1 17syn+ at an MOI of 10 and treated with indicated proteasome inhibitors or DMSO vehicle control. After 8 hours of treatment, total protein was collected and analysed by Western blotting. (A) Expression of RPB1 (red), phosphorylated CTD Ser2 (pS2, green), vinculin, p53, and HSV-1 protein ICP0. Star indicates a high MW band likely corresponding to ubiquitinated RPB1 with arrows indicating accumulated degradation products. (B) Expression of RPB1 (red) and phosphorylated CTD Ser7 (pS7, green) by Western blotting. White lines are the result of cropping irrelevant samples.

**S4 Fig. Quantification of RPB1 levels after stress using different antibodies.** Human foreskin fibroblasts were subjected to heat (44°C, 2h), HSV-1 infection (strain 17syn+, MOI 10, 8h p.i.), high salt (80mM KCl, 1h), and arsenite (0.5mM NaAsO_2_, 1h) treatment and total protein harvested at the end of the stress period. Quantification of RPB1 was performed using the listed antibodies via Western blotting and normalizing to Vinculin levels on the same membrane. Plotted are the means of three biological replicates with standard deviations with representative Western blots to the right.

**S5 Fig. RPB1 serine CTD phosphorylation and viral replication in HSV-1-infected RPB1 K1268R cells.** Clonal cell lines D12 and E2 bearing the RPB1 K1268R mutation and parental wild-type (WT) 293T cells were infected with HSV1(17+)-LoxCheVP26 expressing the VP26 viral late gene fused to mCherry and harvested at 8h p.i. (A) Representative Western blots of CTD serine phosphorylations from data graphed in Fig 3. (B) Viral gene expression at analyzed timepoints is comparable across cell lines as viewed by live-cell imaging of mCherry-VP26 fusion protein. (C) Time-course of HSV-1 strain 17+ production from an initial infection of MOI 0.01. Plotted are the means of two replicates with standard deviations. PFU, <u>p</u>laque-forming <u>u</u>nits; ND, <u>n</u>o plaques <u>d</u>etected for this timepoint; LOD, statistical limit <u>o</u>f <u>d</u>etection defined by 10 plaques at lowest dilution.

**S6 Fig. Comparison of RPB1 levels in the amanitin-based replacement assay.** Cells treated in parallel for the RPB1 replacement assay described in Fig 5 were harvested at the time of infection and probed for total CTD levels, the HA tag from the *trans*-expressed RPB1 constructs, and vinculin as a housekeeping gene.

S1 Table. Antibodies used in this study.

